# *NAC-NOR* mutations in tomato Penjar accessions attenuate multiple metabolic processes and prolong the fruit shelf life

**DOI:** 10.1101/200295

**Authors:** Rakesh Kumar, Vajir Tamboli, Rameshwar Sharma, Yellamaraju Sreelakshmi

## Abstract

Several Penjar accessions of tomato (*Solanum lycopersicum*), widely grown in the Mediterranean region, exhibit prolonged shelf life, and harbor *alcobaca* mutation with valine-106-aspartic acid substitution in the NAC-NOR protein. To uncover the metabolic basis underlying the prolonged shelf life, we compared four Penjar accessions to Ailsa Craig (AC). Three accessions bore *alcobaca* mutation, whereas fourth was a novel *NAC-NOR* allele with only six amino acids in the encoded protein. The cuticle composition among Penjars varied widely during the progression of fruit ripening. All Penjars exhibited delayed ripening, prolonged on-vine and off-vine shelf life, low ethylene emission and carotenoid levels albeit with accession-specific differences. Metabolic profiling revealed shifts in Krebs cycle intermediates, amino acids, and β-aminobutyric acid levels indicating the attenuation of respiration in Penjars during post-harvest storage. The prolonged shelf life of Penjar fruits was associated with a concerted downregulation of a number of cell-wall modifying genes and cell-wall-related metabolites. The accumulation of higher ABA and sucrose levels at the onset of senescence in Penjar fruits likely contribute to reduced water loss. Our analyses reveal that in addition to specialized cuticle composition, the attenuation of various metabolic processes by *NAC-NOR* mutation likely prolongs the shelf life of Penjar fruits.

**Highlight:** The prolonged shelf life of tomato Penjar accessions bearing mutations in NAC-NOR transcription factor appears to be regulated by a combined effect of attenuation of respiration, altered cuticle composition, enhanced ABA and sucrose levels in fruits and downregulation of cell wall modification

## Introduction

Among the factors that influence the economic value of fruits, the post-harvest shelf life is the foremost, as shorter shelf life causes losses during transportation, distribution, and storage. Consistent efforts have been made over last few decades to decipher the molecular-genetic basis underlying the process of fruit ripening and spoilage (Klee and Giovannoni, 2011; Pirrello, 2009; Seymour et al. 2013). Genetic regulation of fruit ripening is mainly deciphered in tomato, where spontaneous mutants such as *rin*, *nor* and *Cnr*, fail to undergo characteristic accumulation of lycopene in the fruits and remain firm for a long time (Giovannoni, 2007). The *RIN*, *CNR* and *NOR* genes encode for transcription factors belonging to MADS, SBP box, and NAC family respectively (Seymour et al. 2013).

The *nor* mutant family has two alleles; *nor* with truncated protein, and *alcobaca (alc)* with a single amino acid substitution (Casals et al. 2012; Giovannoni, 2004). The cultivars harboring either of these alleles show prolonged shelf life of fruits, albeit duration depends on the fruit size and genetic background (Garg et al. 2008a,b; Casals et al. 2012; Mutschler et al. 1988). The biochemical characterization of *nor/alc* mutants revealed that compared to normal ripening cultivars, the *nor/alc* fruits had altered cuticle composition with denser cutin matrix (Kosma et al. 2010; Saladié et al. 2007), were deficient in carotenoids accumulation, and were resistant to infections (Garg et al. 2008a,b). As *NOR* mutation is strongly associated with long shelf life, search for additional NAC family transcription factors revealed a role for *NAC1* and *NAC4* in tomato ripening (Ma et al. 2014; Zhu et al. 2014).

The onset of ripening in tomato initiates several irreversible processes such as changes in metabolite composition, loss of chlorophyll, accumulation of carotenoids and cell wall softening (Carrari et al. 2006; Fraser et al. 2007; Giovannoni, 2007). These responses are triggered by the climacteric rise of ethylene in the fruits, a process that is subdued in the *nor* mutants. Current evidences indicate that in addition to ethylene, other hormones like abscisic acid (ABA), jasmonic acid (JA), methyl jasmonate (MeJA), auxin, etc. also affect fruit ripening and quality (Kumar et al. 2014; McAtee et al. 2013). It is believed that *NOR* acts upstream to ethylene in regulating the ripening of tomato (Giovannoni, 2007). However, it is not known whether *NOR* influences the levels of other hormones during ripening.

The studies using transgenic manipulation of genes revealed that softening of tomato fruits can be prevented by silencing of ethylene biosynthesis (Oeller et al. 1991; Xie et al. 2006) and perception (Tieman et al. 2000), increasing polyamine biosynthesis (Nambeesan et al. 2010), suppression of ABA biosynthesis (Sun et al. 2012), and downregulation of enzymes participating in cell wall degradation (Brummell et al. 2002; Vicente et al. 2007; Uluisik et al. 2016). The transpirational water loss through cuticle also affects the fruit firmness. Thus fruits that retain cellular turgidity remain firm for longer periods (Saladié et al. 2007). From the foregoing, it is apparent that a combination of multiple processes likely regulates prolonged shelf life of *nor/alc* mutants. However, only limited information is available about modulation of these processes in the *nor/alc* mutants (Casals et al. 2015; Saladie et al. 2007, Osorio et al. 2011).

Breeders have used *alc* for better fruit quality, though hybrid fruits have a slightly shorter shelf life than *rin* or *nor* hybrids (Garg et al. 2008a,b). Among the landraces grown in Mediterranean, the Penjars are popular for long post-harvest shelf life. Several Penjar accessions bear the *alc* mutation (Casals et al. 2012), however, little is known about the metabolic profiles of fruits during postharvest storage. In the current study, we analyzed four Penjar accessions during ripening and post-harvest storage to decipher the metabolic basis for the prolonged shelf life.

## Materials and Methods

### Plant material and post-harvest analysis

Penjar accessions (a gift from RF Muñoz) and tomato (*Solanum lycopersicum*) cultivar Ailsa Craig were grown in pots between October to February (day 30±2°C, night 18±2°C) season, which is similar to Mediterranean summer. The fruits were collected at mature green (MG), breaker (BR) and red ripe (RR) stages. For Penjars, attainment of uniform fruit coloration was considered equivalent to RR stage. The pericarps were snap-frozen in liquid nitrogen and stored at −80°C until analysis. Freshly harvested RR fruits were incubated at 24±2°C under natural day/night conditions. The appearance of wrinkling was considered as the onset of senescence (SEN). On post-harvest storage, the Penjar-2 fruit was first to show wrinkling at 65-days, which was considered as SEN. The weight loss during post-harvest storage was monitored by periodic weighing of fruits.

### Sequence, SNP, promoter and SIFT analysis

The *NOR* promoter (1834 bp) and cDNA were amplified from AC and Penjars (Primer details in Table S1), and PCR products were sequenced (Macrogen, South Korea). The SNPs were identified using Multialin Interface page (Corpet et al. 1988). The amino acid sequences were aligned using Multialin or PRALINE software (http://www.ibi.vu.nl/programs/pralinewww/) (Bawono and Heringa, 2014). The promoter sequences (3 Kb) of genes were retrieved from SOL genomics network (https://solgenomics.net/) and analyzed for NAC transcription factor binding sites using PlantPAN 2.0 (http://PlantPAN2.itps.ncku.edu.tw) (Chow et al. 2016). The deleterious effect of non-synonymous polymorphisms was calculated by SIFT (www.sift.dna.org; version 4.0.5) (Sim et al. 2012). For SIFT score ≥0.05, SNPs were predicted as tolerated, whereas score ≤ 0.05 was predicted to be deleterious for the protein function.

### Carotenoids, hormones, transcripts and fruit metabolites analysis

The carotenoid content was estimated using Gupta et al. (2015) protocol. The ethylene emission and hormone analysis were carried out as described in Kilambi et al. (2013) and Bodanapu et al. (2016) respectively. RNA was extracted from fruits in three biological replicates using hot phenol method (Verwoerd et al. 1989) and q-PCR was carried out as described earlier (Kilambi et al. 2013) (Primer details in Table S1). Tomato fruit metabolites were extracted, analyzed and identified using GC-MS as described in Bodanapu et al. 2016.

### Analysis of cuticle components

For cuticular wax and cutin monomers analysis, five biological replicates at MG and RR were used. The total cuticular wax was isolated as described by Leide et al. (2007). Intact fruits were dipped for 2 min in chloroform and 5-Pregnen-3β-ol-20-one was added as an internal standard. The solvent was evaporated under nitrogen to 1 mL followed by drying in a Speed Vac before GC-MS. The cuticle membrane (CM) of fruits was isolated using pectinase and cellulase in 50 mM citrate buffer (pH 4.5, supplemented with NaN_3_). Isolated 1 cm^2^ size CMs were repeatedly washed with MilliQ water and air dried. CMs were dewaxed as described by Kosma et al. (2010). Dried CMs were treated with chloroform: methanol (1:1; v/v) for 24 h followed by washing with methanol (4-5 times) to remove chloroform. The dewaxed CMs were depolymerized by alkaline hydrolysis (Osman et al. 1995) and derivatized for 30 min with 80 μL MSTFA at 37°C. 10 μL of 5-Pregnen-3β-ol-20-one (1 mg/mL) was used as internal standard for cuticular wax and cutin analysis.

For GC-MS, 1 μL sample was injected to RXi column as described for fruit metabolites in Bodanapu et al. 2016 in a splitless mode. Cutin and wax components were separated using the following program: 1 min at 50°C with a linear ramp of 10°C/min to 180°C and held at 180°C for 2 min, again a linear ramp of 3°C/min to 300°C and then held at 300°C for 18 min. Both ion source and injector temperature were 250°C and helium was used as carrier gas at a flow rate of 1.5 mL/min. The mass spectra were recorded at a scan rate of 2 scans/sec with a scanning range of 40 to 850 m/z. The raw data were processed, and metabolite identity was assigned as described above for GC-MS analysis.

### Statistical analysis

For all experiments, a minimum of 3-5 biological replicates was used and mean with the standard error was calculated. The **StatisticalAnalysisOnMicrosoft-Excel** software (http://prime.psc.riken.jp-/Metabolomics_Software/StatisticalAnalysisOn-MicrosoftExcel/) was used to obtain significant differences between data points using Student’s test (P ≤ 0.05). PCA was carried out using metaboanalyst 3.0.

### Network analysis

The carotenoid, hormone, *PSY1* and *CYCB* abundances for Penjars and AC were pair-wise correlated at MG, BR and RR stages (n=3, P-value ≤ 0.05) using the Pearson’s Correlation coefficient (PCC). The associations with a PCC value (*r*) ≥ 0.8 (+/-) were used to create a network, and were visualized using Cytoscape (http://www.cytoscape.org/) (Shannon et al. 2003). The red and gray edges indicate negative and positive correlations respectively.

### Accession Numbers

The accession numbers of genes examined in this study are given in Table S1 along with the primer sequences used for amplification.

## Results

Penjar tomatoes cultivated in northeastern Spain are harvested during July to September for consumption in winter owing to their longer shelf life. In this study, we compared the cuticle composition, primary metabolites, hormones, carotenoids composition, and cell wall modification in four Penjar accessions with a normal ripening cultivar, Alisa Craig (AC).

### **Penjar-1 has two novel *NOR* mutations**

To uncover the genetic basis of prolonged shelf life, we amplified and sequenced the full-length cDNA of *NOR* gene from all four Penjars (Figure 1; Fig. S1A; Table S1). Notably, two mutations present in Penjar-1 were novel, the C to A transversion at 20^th^ position created a stop codon resulting in a truncated NOR protein with only six amino acids (Figure 1) and downstream C to A transversion at 37^th^ position led to glutamine to lysine substitution at the 13^th^ position of the protein. Excepting Penjar-1, other three accessions possessed a single base change similar to *alc* mutant (T to A at 316^th^ position of cDNA; substitution of valine to aspartic acid at 106^th^ position of NOR protein, Fig. S1B) akin to the mutation present in 27 Penjar accessions (Casals et al. 2012). The low SIFT score (<0.05) (Ng and Henikoff, 2003; Sim et al. 2012) predicted that the mutations in all four accessions are deleterious for NOR protein function. The examination of *NOR* gene promoter (1834 bp) did not reveal any additional SNPs in above accessions (Fig. S1C).

**Figure 1.**
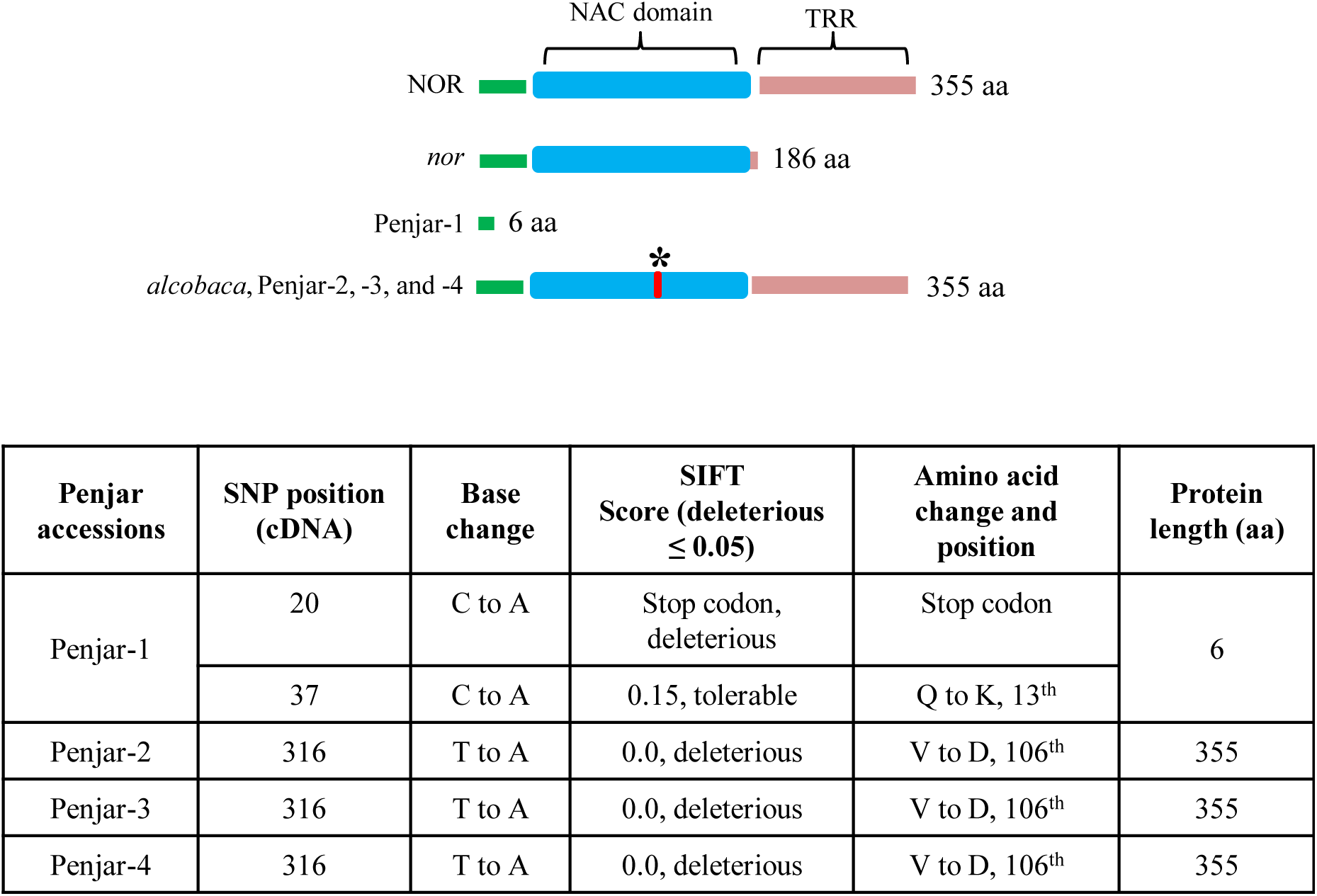
Mutations in *NAC-NOR* gene in *nor, alcobaca* and Penjar accessions. AC encodes a full-length NOR protein of 355 aa. *nor* mutant produces a truncated protein of 186 aa. The Penjar-1 encodes a protein of 6 aa length, whereas *alc*, Penjar-2, -3, and -4 encode a full-length protein with a single amino acid change of V to D at 106th position (indicated by an asterisk and red band). NAC indicates NAC domain and TRR indicates transcriptional regulatory region. The details of base changes and the corresponding positions in the cDNA, protein, and SIFT score are indicated in the Table.

### Penjars exhibit delayed ripening and prolonged shelf life

The tomato fruits are considered fully ripe when the fruits acquire uniform red coloration, albeit ripe Penjar fruits exhibited different colors. At the ripe stage, fruits of Penjar-2 were light red, Penjar-1 were yellow-orange and Penjar-3 and -4 were orange colored (Fig. S2A). The attainment of uniform coloration by Penjar fruits was considered equivalent to the *r*ed *r*ipe stage (RR) of AC.

The fruit growth of Penjars was nearly similar to AC up to *m*ature *g*reen stage (MG). Post-MG stage, the transition period to *br*eaker stage (BR) in Penjars and AC was nearly similar and also akin to *nor, alc* and DFD (Mutschler et al. 1992; Saladié et al. 2007; Simpson et al. 1976) (Fig. S2B). However, BR to RR transition (8-15 days) was more prolonged in Penjars than AC, with Penjar-1 being slowest (15 days). Apart from delayed ripening, all Penjars exhibited extended shelf life both on-vine and off-vine. The on-vine ripened Penjar fruits manifested no visible signs of shriveling even after 130 days *p*ost-*a*nthesis (DPA) (Simpson et al. 1976), while AC fruits shriveled at 70-80 DPA (Fig. S3).

We next examined in detail the off-vine shelf life of Penjars and AC. The fruits harvested at RR were monitored for wrinkling at regular intervals. In AC, wrinkling appeared on 10^th^-day post-harvest (DPH), became prominent by 20 DPH followed by shrinking of fruits due to water loss (Figure 2A). At 65 DPH, the shrinkage was markedly apparent, and fruits lost 40% of initial weight. Contrastingly, all Penjar fruits even after 65 DPH retained 80-90% of initial weight and displayed no wrinkling (Figure 2B). By 80 DPH, AC fruits were completely shriveled, whereas, Penjar-1, -3 and -4 showed no signs of wrinkling even at 150 DPH. Among these, Penjar-2 fruits manifested wrinkling by 65 DPH; therefore 65 DPH was selected as the onset of senescence (SEN) for the subsequent post-harvest studies.

**Figure 2.**
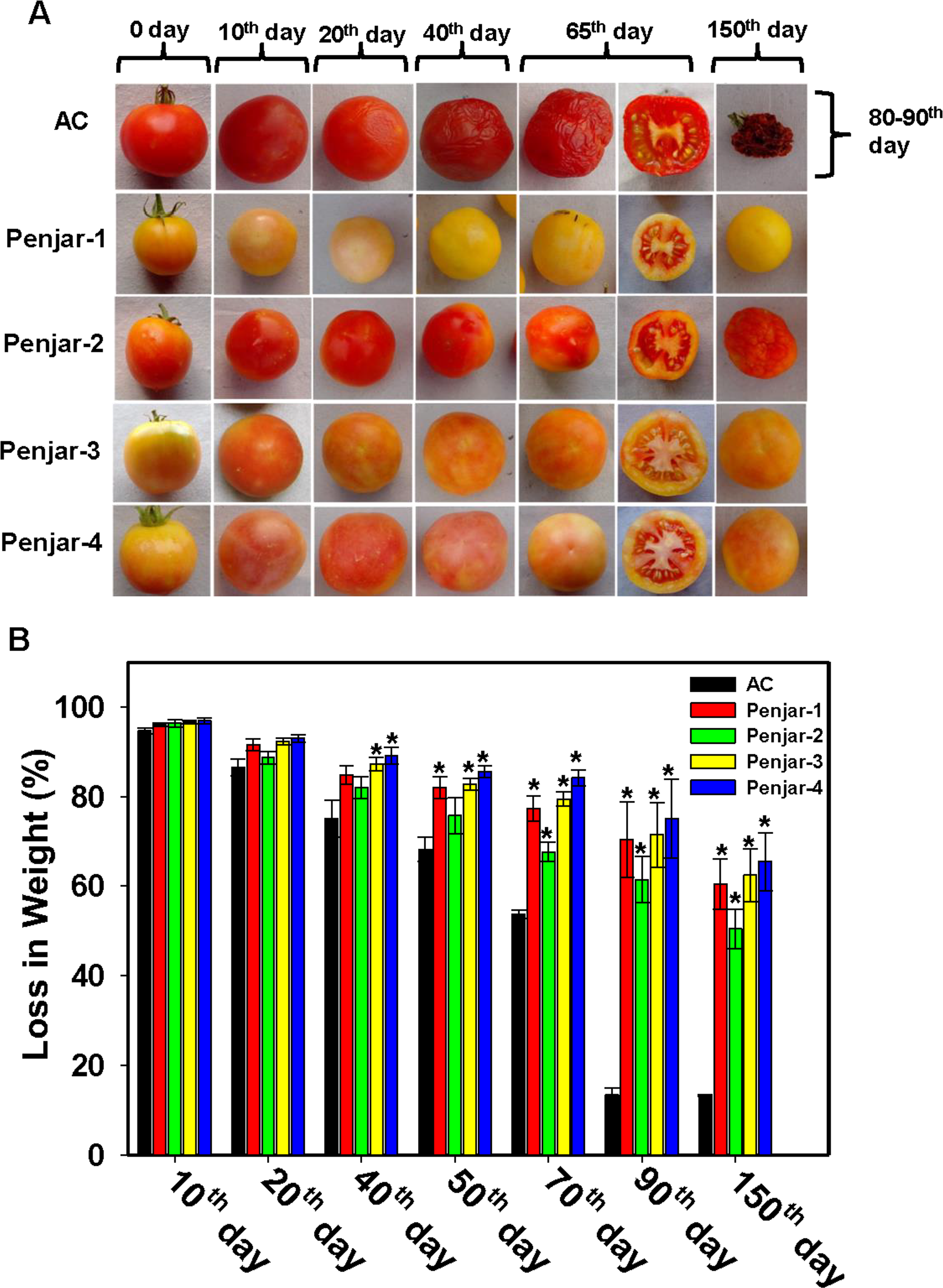
Post-harvest shelf life and water loss in AC and Penjar fruits. A, The fruits were harvested at ripe stage and incubated under normal day-night conditions as described in methods. By 40 days post-harvest, AC fruits showed prominent wrinkling, and 65 day-old fruits displayed a clear sign of cellular disintegration. **B**, The water loss was measured during post-harvest storage. Data are means of 5 biological replicates ± SE, ‘**\***’ indicates P ≤ 0.05.

### ***NOR* and *RIN* expression is altered in Penjars**

In tomato, *RIN* and *NOR* are two key regulators of ripening and shelf life (Giovannoni, 2001). To ascertain the role of these regulators in ripening and shelf life of Penjar fruits, their transcript levels were analyzed by qRT-PCR (Fig. S4). Consistent with the long shelf life, the *NOR* expression in Penjar fruits at RR was significantly lower than AC. The *NOR* expression declined in AC and Penjars (except Penjar-1, -2) at SEN. Interestingly, *RIN* transcript was below the detection limit at MG in AC and thereafter increased progressively. The reduced expression of *RIN* in Penjar-1 at RR and SEN is in consonance with it being a knockout mutant. Other Penjars showed higher *RIN* expression at RR and the levels were similar to AC at SEN.

### Penjars show wide variation in cuticle composition

The cuticle protects fruit firmness by preventing transpirational water loss. Consistent with this, stripping of the cuticular wax (CW) by chloroform accelerated shrinkage (Figure 3A) and water loss of Penjar fruits akin to AC (Fig. S5A). The analysis of CW revealed the presence of nearly 100 compounds that were classified into five categories: hydrocarbons (alkanes, alkenes), fatty acids, fatty acid alcohols, aromatic and miscellaneous compounds (sterols and triterpenoids) (Fig. S5B, Table S6). At MG, the CW levels were very high in Penjars than AC barring Penjar-1. At RR though CW content increased in all Penjars and AC, only Penjar-3, -4 had levels higher than AC (Figure 3B).

**Figure 3.**
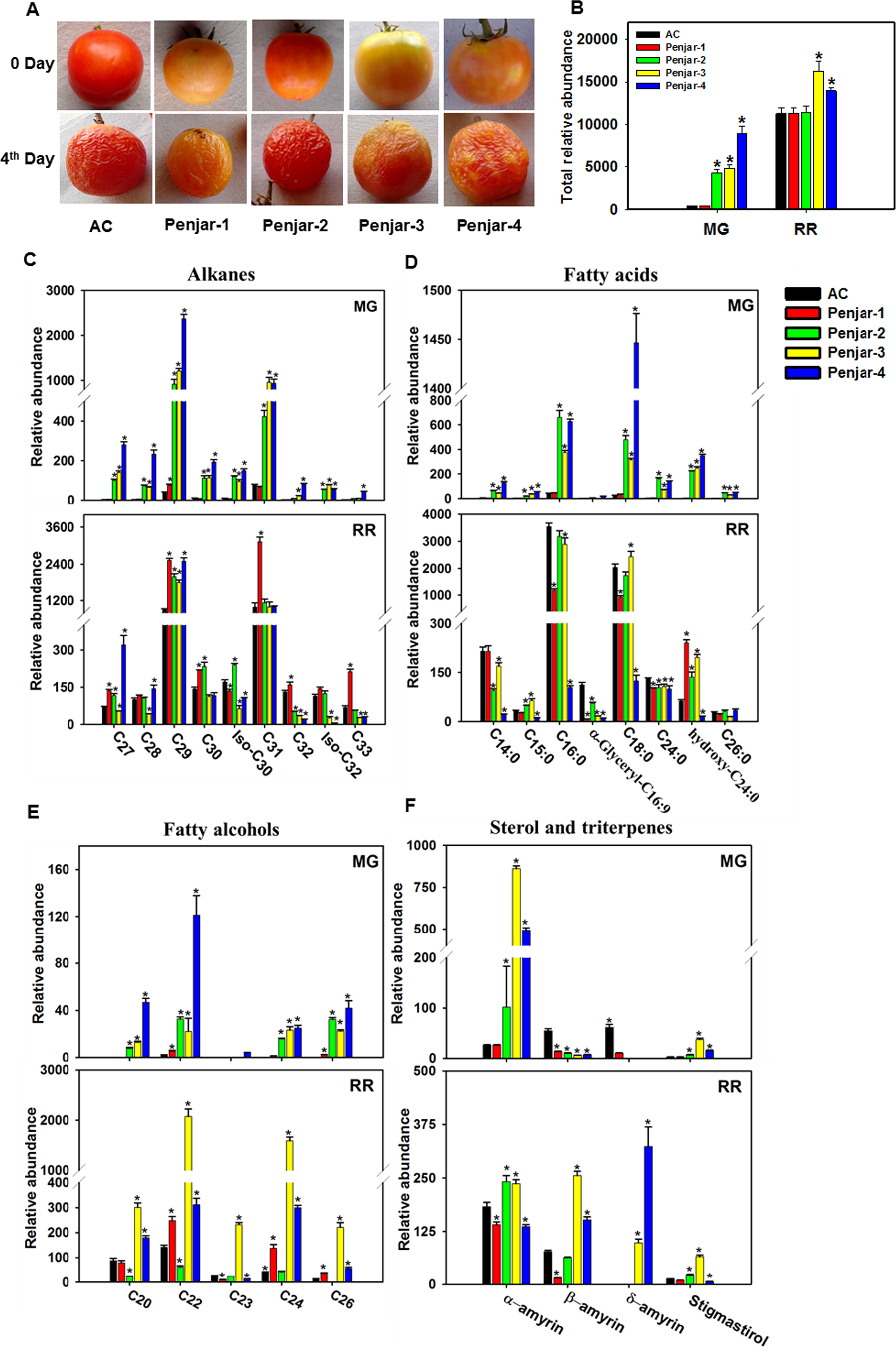
The cuticular wax composition of AC and Penjar fruits during ripening. **A**, The fruits were harvested at the ripe stage, and cuticular wax from the surface of both AC and Penjar fruits was removed by using chloroform. This resulted in the severe shrinking of fruits due to loss of moisture. **B**, The total cuticular wax content of AC and Penjar fruits at MG and RR. **C-F**, The relative abundance of cuticular wax components: alkanes (**C**), fatty acids (**D**), fatty alcohols (**E**), sterols and triterpenes (**F**) in MG and RR fruits of AC and Penjars. Data are means of 3 biological replicates ± SE, ‘**\***’ indicates P ≤ 0.05.

The relative abundance of different cuticle components distinctly differed between four Penjars as well as with AC (Figure 3B-F; Fig. S5). The abundance of nine detected alkanes in RR fruits widely differed in Penjars and AC, excepting C29 alkane that was 1.5-2 fold higher in Penjars than AC. While few fatty acids classes at MG were more abundant in Penjars than AC, the abundance of these fatty acids distinctly changed at RR. Several fatty acid alcohols were most abundant in Penjars particularly at MG, and their levels significantly increased during ripening. Compared to stigmasterol, the levels of all triterpenes changed during the transition to RR, albeit variably among different Penjars and AC.

Several cutin components such as glycerol, hydroxy hexadecanoic acid, octadecanoic acid, 18-OH octadecanoic and 9,18-diOH octadecanoic were higher in all four Penjars at MG compared to AC (Table 1). However, at RR only glycerol was higher in all four Penjars than AC. The other cutin components in Penjars though varied did not show any concerted increase or decrease like glycerol on comparison to AC. Analogously, the analysis of transcript levels of fourteen genes putatively regulating cuticle biogenesis showed upregulation of *cutin deficient 2* (*CD2)*, *MIXTA*, *ABC transporter G family member 11* (*WBC11)*, *glycerol-3-phosphate acyltransferase 6* (*GPAT6)*, *3-ketoacyl-CoA reductase 1* (*KCR1)*, *enoyl-CoA reductase (ECR)*, and downregulation of *fatty acyl-ACP thioesterase* (*FATB)* in MG fruits of all Penjars compared to AC (Fig. S6; Table S1). However, at RR only *FATB* transcript was commonly altered reinforcing that the cuticle biogenesis is more specifically modulated in Penjars at MG than at RR.

**Table 1.**
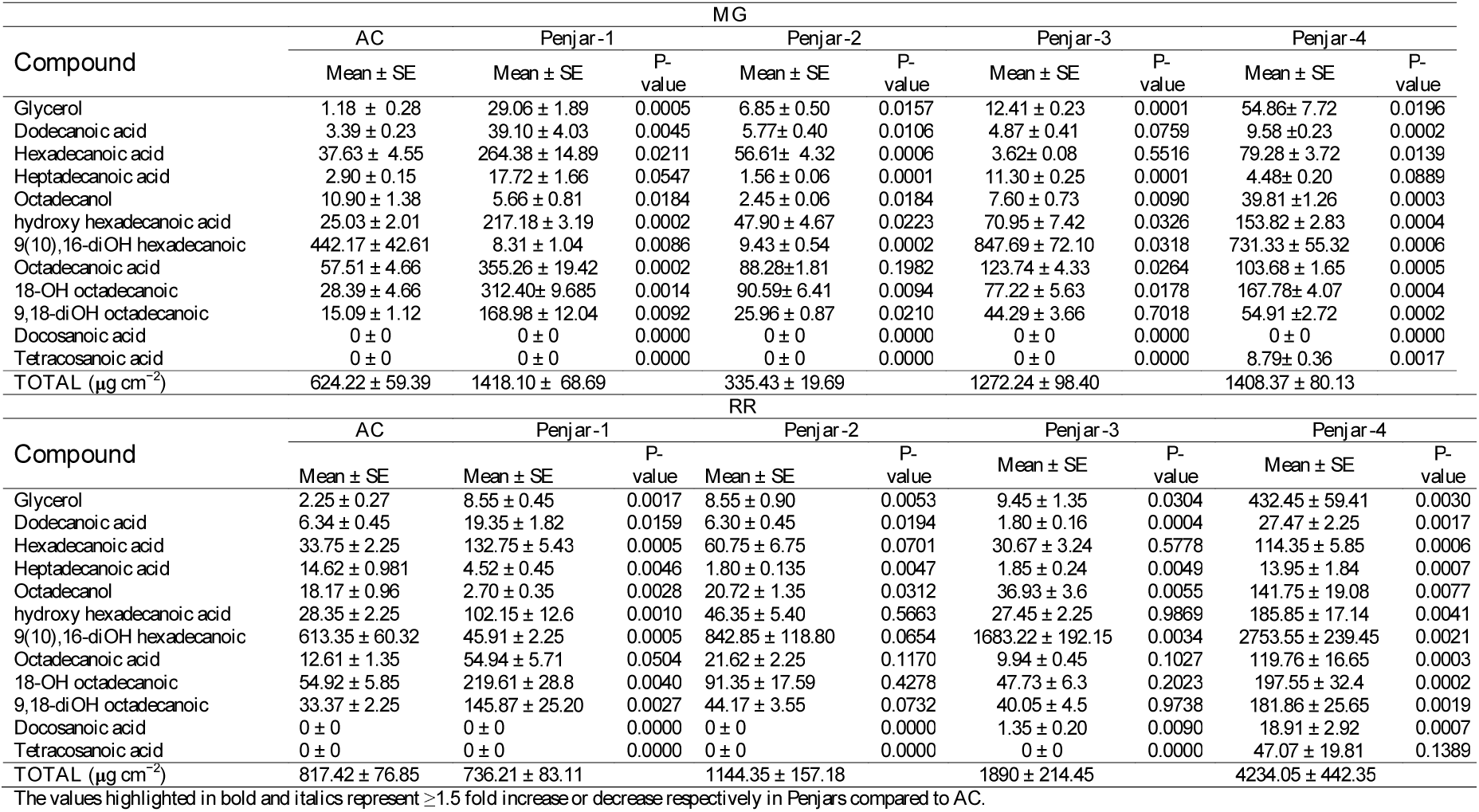
List of compounds identified in the cutin matrix of MG and RR stage fruits of AC and Penjars.

### Cell wall modifying genes are downregulated in Penjars

In tomato, fruit ripening is accompanied by progressive softening of cell walls by a battery of cell-wall specific enzymes, whose suppression extends fruit shelf life (Brummell et al. 2002; Cantu et al. 2007; Meli et al. 2010; Uluisik et al. 2016). To ascertain whether the extended shelf life of Penjar fruits is linked with the reduced expression of genes encoding cell wall modifying enzymes, expression of *polygalacturonase* (*PG-2A, PG-β), expansin (EXP), pectin methylesterase (PME), galactosidase (α-GAL, β-GAL), mannosidase (α-MAN, β-MAN), deoxyhypusine synthase (DHS)* and *hexosaminidase* (*HEX*) was examined. Consistent with the above notion, expression of at least eight cell wall modifying genes- *β- GAL*, *α-GAL*, *α-MAN*, *β-MAN*, *PG-*β, *PME*, *EXP*, and *DHS* was downregulated at RR of Penjar fruits (Figure 4). Only *PG2A* and *HEX* genes showed higher expression at RR in Penjar-1, and -2 than AC. At SEN, Penjars showed accession-specific variation in gene expressions. All Penjars showed higher expression of *α-MAN*, *β-MAN*, *PG-*β genes than AC. Among different Penjars, expression of *α-GAL* and *PG2A* was greater in Penjar-1, and -3, *β- GAL* was elevated in Penjar 2, *PME* and *EXP* was higher in Penjar-3, and *DHS*, *HEX* was enhanced in Penjar-3, -4 than AC. The analysis of promoters of above genes revealed the presence of multiple NAC binding sites (Fig. S7; Table S4).

**Figure 4.**
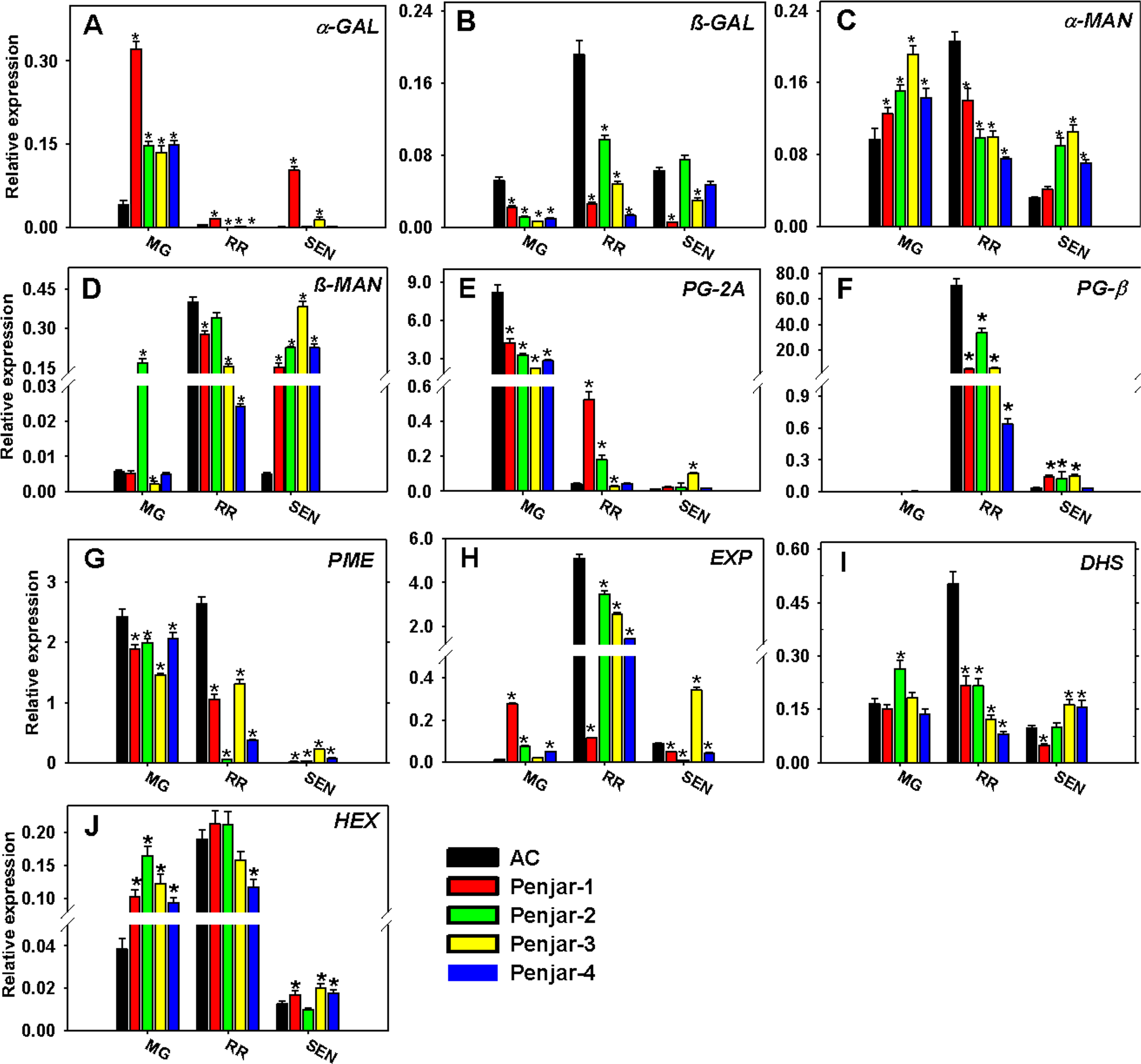
Relative expression of cell wall modifying genes in AC and Penjar fruits. The fruits were collected at different ripening and SEN stages, and expression of various cell wall modifying genes was determined by qPCR. The graphs represent the data obtained after normalization with *actin* and *ubiquitin*. **A**, *α-Galactosidase;* **B**, *β-Galactosidase;* **C**, *α- Mannosidase;* **D**, *β-Mannosidase;* **E**, *Polygalacturonase 2A;* **F**, *Polygalacturonase β-subunit;* **G**, *Pectin methylesterase;* **H**, *Expansin;* **I**, *Deoxyhypusine synthase* and **J**, *β-N-acetyl-D- hexosaminidase*. Data are means of 3 biological replicates ± SE, ‘**\***’ indicates P ≤ 0.05.

### Hormone profiling

In tomato, a climacteric fruit, ethylene production is coupled to ripening and the associated fruit coloration and softening (Alexander and Grierson, 2002). Though the overall pattern of ethylene emission was similar in AC and Penjars, consistent with delayed ripening and lower carotenoid content, the Penjar fruits emitted significantly less ethylene than AC (Fig. S8A), excepting Penjar-2 that emitted slightly higher ethylene at BR. In consonance with reduced ethylene emission, at RR, the transcript levels of key genes of system II ethylene biosynthesis - *1-aminocyclopropane-1-carboxylate synthase 2 and 4 (ACS2, ACS4)*, and at BR of *1-aminocyclopropane-1-carboxylic acid oxidase 1 (ACO1)* were significantly lower in Penjars than in AC (Fig. S8B-D). In contrast, system I ethylene biosynthesis gene-*ACO3* levels (Fig. S8E) were similar in Penjars and AC at BR (except Penjar-3 and -4) and RR (except Penjar-2 and -4). Nearly equal gene expression of *ACS2*, *4* and *ACO1* at BR in Penjar-2 and AC may be related to slightly higher ethylene emission from Penjar-2 fruits.

In tomato, other hormones like jasmonate (JA), methyl jasmonate (MeJA) and salicylic acid (SA) also play important roles in lycopene accumulation (Kumar et al. 2014; Liu et al. 2012). In both AC and Penjars, the transition to RR upregulated JA levels. However, the stimulation was substantially lower in Penjars (Figure 5). The upregulation of JA was sustained at SEN in AC, and light-red fruited Penjar-2, while it declined in yellow/orange-fruited Penjars. Contrastingly, Penjar accessions showed changes in MeJA levels that were opposite to AC during the transition from MG to BR with a steady level at RR (except Penjar-2) and a decline at SEN. In tomato, lycopene accumulation in fruits requires higher SA levels during early stages of ripening (Ding and Wang, 2003). Consistent with this, while SA level in MG/BR fruits of AC was significantly higher than RR, it was much lower in Penjars at MG and BR.

**Figure 5.**
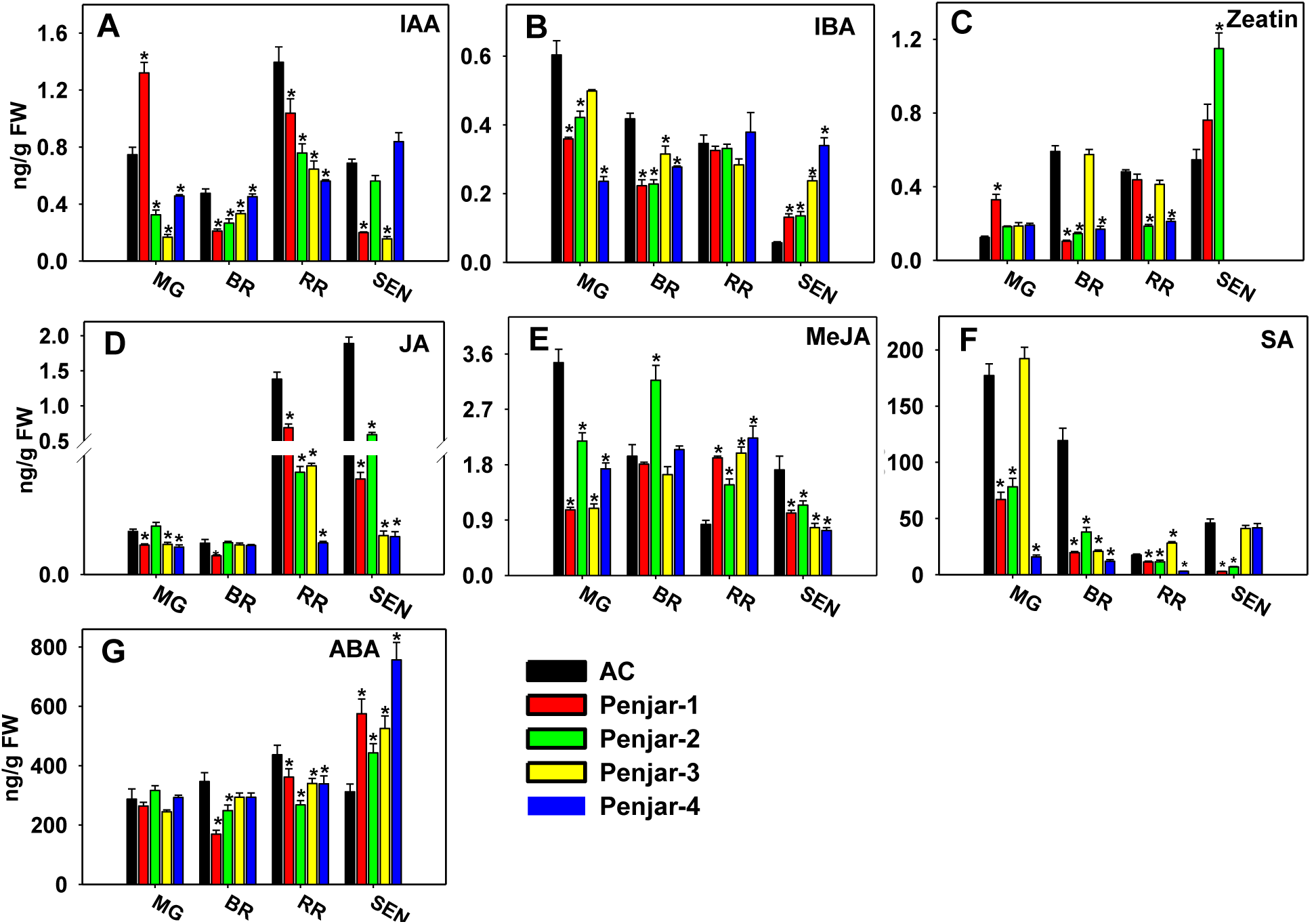
Hormone profiling in AC and Penjar fruits. The fruits were collected at different ripening and SEN stages, and hormone profiles were determined as described in methods. **A**, IAA; **B**, IBA; **C**, zeatin; **D**, JA; **E**, MeJA; **F**, SA; **G**, ABA. Data are means of 5 biological replicates ± SE, ‘**\***’ indicates P ≤ 0.05.

Also, the levels of abscisic acid (ABA), indole-3-acetic acid (IAA), indole-3-butyric acid (IBA) and zeatin were altered in Penjar fruits (Figure 5). In AC, ABA level increased during ripening and then declined at SEN, whereas Penjars showed only modest variations during ripening, but with upregulation at SEN. Both Penjars and AC did not show a consistent pattern for IAA levels, except that Penjars had significantly low IAA levels (barring Penjar-1 at MG). Conversely, the IBA levels gradually declined in AC from MG to SEN, and Penjar-4 retained consistently steady IBA levels at all stages. In higher plants leaves, cytokinin reportedly inhibits the senescence process (Lim et al. 2007), however, in both AC and Penjars, zeatin levels did not correlate with fruit senescence.

### **Penjars show reduced *PSY1* expression**

Since ripe Penjar fruits do not acquire typical deep red coloration, it was assumed that *NOR* mutations might have influenced the carotenogenesis in fruits. Profiling of carotenoids at different stages revealed substantially low carotenoid levels in Penjars (Fig. S9), akin to *nor* and *alc* (Kopeliovitch et al. 1979; Sink Jr. et al. 1974). Consistent with light-red color, Penjar-2 fruits accumulated higher levels of phytoene, phytofluene, and lycopene than other three Penjars. In tomato, *phytoene synthase 1* (*PSY1*) and chromoplast-specific *lycopene β- cyclase* (*CYCB*) genes are two key regulators of carotenogenesis in fruits, and their increased expression from MG to RR is closely associated with carotenoid accumulation (Hirschberg, 2001). The reduced *PSY1* expression at RR is consistent with lower carotenoid levels in respective Penjars, with the least reduction in Penjar-2 (Fig. S10). Interestingly, Penjar-1 fruits showed 2-fold higher *CYCB* expression at RR than AC, which is reflected as high β-carotene/lycopene ratio than other Penjars (Table S2A). The transition from RR to SEN in Penjar fruits was marked by an accelerated loss in carotenoids levels (∼50-70%) than AC (26%) (Table S2B). Among *PSY1* and *CYCB*, at SEN, the *PSY1* expression declined more severely in Penjars than in AC. In contrast, *CYCB* expression though decreased in Penjars yet was 2-3 folds higher than AC. The correlation networks (*r* = 0.8) consisting of most abundant carotenoids, *PSY1* and *CYCB* expression and the hormones during ripening revealed interesting patterns (Fig. S11; Table S3). Penjar-1, -2 -3, and AC showed a strong positive correlation between JA and carotenoid levels, and ethylene and *PSY1* expression levels. In contrast, Penjar-4 showed very few interactions compared to other Penjars and AC. Though other hormones also interacted with different carotenoids and transcripts, these varied between different Penjars.

### Metabolite analysis in Penjars

Using GC-MS, we identified ∼110 primary metabolites in the MG, BR, RR and SEN fruits of AC and Penjars (Table S5). Notwithstanding the diverse metabolite composition in AC and different Penjars, principal component analysis (PCA) revealed that the metabolite profiles of ripening Penjars were closer and overlapped with each other (Fig. S12A). At SEN, while the profiles were closer in PC1, they showed accession-specific differences in PC2 (Fig. S12B). Based on functional groups, the identified metabolites were classified as organic acids, amino acids, amines, fatty acids, and sugars. On the metabolic pathway, only those metabolites with ≥1.5 fold (Log_2_ Penjar/AC value of 0.58; P ≤0.05) upregulation or downregulation in Penjar fruits compared to AC were mapped (Figure 6).

**Figure 6.**
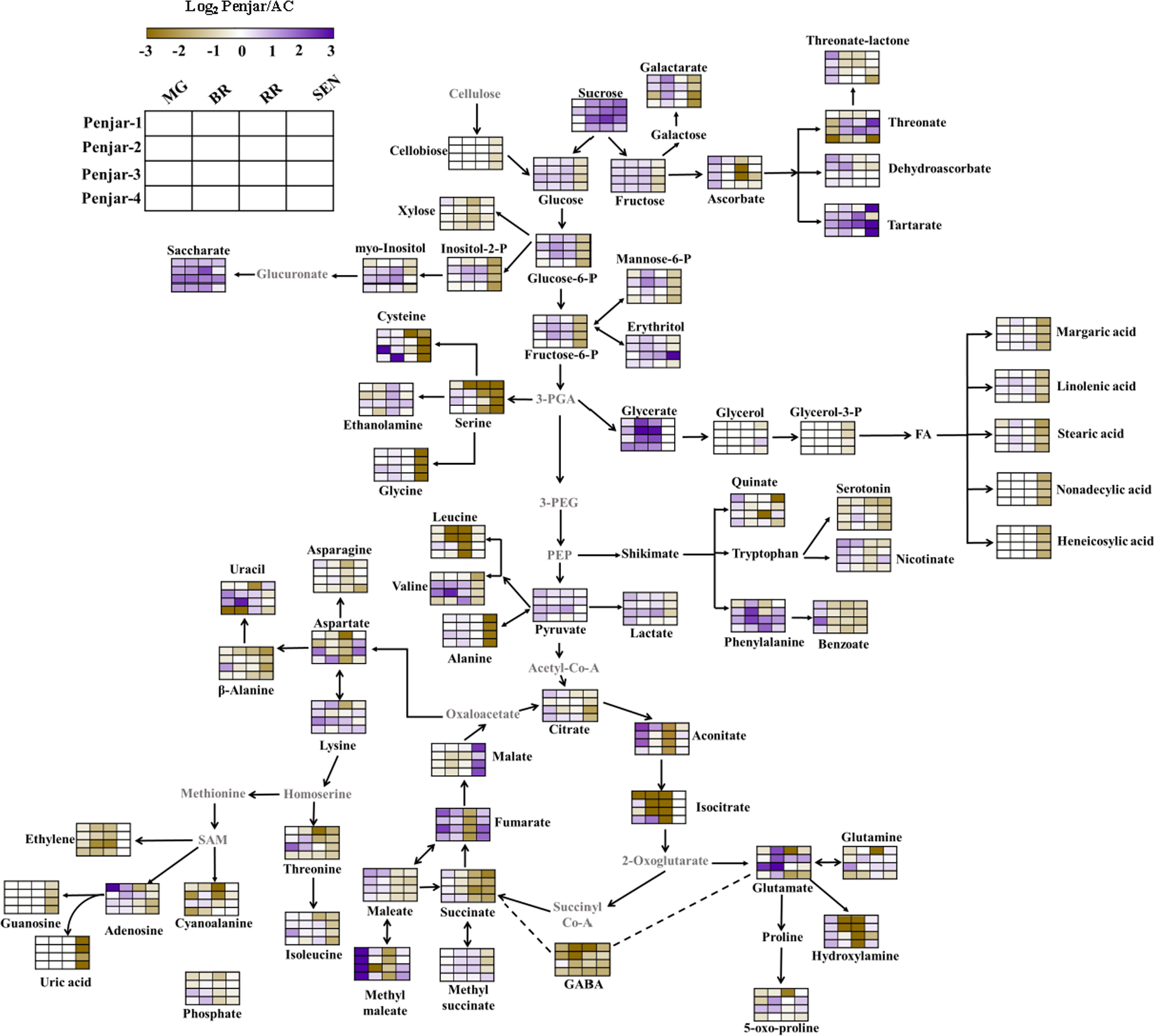
The metabolic shifts in the fruits of Penjars in comparison to AC during ripening and post-harvest storage. The relative abundances of the metabolites in Penjar fruits at MG, BR, RR and SEN stages were obtained from the Log_2_ Penjar/AC values, and only those metabolites with ≥1.5 fold change (n=5 ± SE, P ≤0.05) were mapped on the pathway. The scale at the top left corner represents Log_2_ fold changes in the range of -3 to +3. The metabolites indicated in gray letters on the pathway were not detected in the GC-MS analysis.

### Penjars show differential accumulation of Krebs cycle intermediates

The fruits of Penjars displayed differential accumulation of Krebs cycle intermediates during ripening and postharvest storage (Fig. S13). Particularly in Penjars, organic acids levels were lower than AC during ripening. Citrate, the most abundant organic acid in ripe fruits, was significantly lower in Penjar fruits at SEN, whereas, at RR, the citrate level was lower in Penjar-1 and -3 than AC. Likewise, the acotinate levels in RR fruits were also highly reduced than AC, albeit at SEN, the levels were nearly similar. Though the levels were low at RR, malate, fumarate and methyl maleate were significantly upregulated at SEN in Penjars. The levels of succinate in Penjar fruits decreased during ripening and SEN.

### Penjars exhibit low levels of amino acids during ripening and post-harvest storage

During tomato ripening, suppression of climacteric rise of ethylene reportedly reduces the amino acid content (Gao et al. 2007). Consistent with reduced ethylene emission in Penjar fruits, the levels of several amino acids (12) were lower than in AC. At MG, free amino acids such as alanine, leucine, isoleucine, valine, glycine, serine, alanine-3-cyano, threonine, aspartic acid, beta-alanine, glutamine, GABA, etc., were relatively low in both AC and Penjar (except Penjar-3) fruits (Fig. S14). The onset of ripening stimulated a modest increase in the amino acids and amines in AC and Penjar-2 fruits at BR. Interestingly, the GABA levels declined in all Penjars by 2.5-5 fold at BR. At RR, in most Penjars, the amino acids such as beta-alanine, GABA, isoleucine, glycine, serine, alanine-cyano, threonine, aspartic acid, asparagine, and hydroxylamine were much lower than AC. At SEN, reduction in the amino acids was most severe in Penjars with ∼10-30 fold decrease in alanine, valine, leucine, and isoleucine. The levels of only a few amino acids such as glutamate at RR of Penjar-2, phenylalanine at RR and SEN of Penjar-3 and -4, and aspartate at SEN of Penjar-2 and -3 were higher than AC at respective stages. All Penjar fruits had reduced level of serotonin (5- hydroxytryptamine) at SEN while serotonin levels were high in AC fruits throughout ripening and SEN.

### Penjars exhibit differential accumulation of sugars and cell wall-related metabolites

During fruit ripening, glucose and fructose, cell wall-derived sugars, and polyols increase, while sucrose and sugar phosphates and fatty acids decrease (Fraser et al. 2007). Ripening Penjar fruits characteristically exhibited substantially higher levels of glucose, fructose, glucose-6-P, sucrose, etc. than AC (Fig. S15). At SEN glucose and fructose levels declined in Penjars but increased in AC. The Penjar fruits at SEN also showed increased levels of glucose-6-P, fructose-6-P, and mannose-6-P. At SEN, high levels of cell wall-related metabolites like xylose and cellobiose in AC fruits and reduced levels of galacturonate and galactarate levels was observed in Penjar fruits.

The profiles of other metabolites such as fatty acids, nucleotides, dehydroascorbate, nicotinate, and quinate in Penjar and AC fruits were also distinctly different (Fig. S16). Though the levels of fatty acids such as margaric acid, linolenic acid, stearic acid in Penjar fruits were similar to AC during ripening, at SEN their levels were higher in AC. The metabolites-nonadecyclic acid and heneicosylic acid that are detected only at SEN were much lower in Penjars than AC. In contrast to Penjars, AC at SEN exhibited increased levels of adenosine and guanosine levels. Interestingly, uracil levels were high in Penjars throughout ripening including SEN except for Penjar-2 that had low levels at SEN.

## Discussion

Tomato being a perishable fruit is the subject of intensive investigations to extend its post-harvest shelf life (Pech et al. 2013). Among the regulatory factors identified, the *nor* and *rin* alleles in tomato prolong the shelf life of fruits, even in heterozygous condition (Garg et al. 2008a; Giovannoni, 2007). The *NOR* gene belongs to NAC transcription factor family (Klee and Giovannoni, 2011; Martel et al. 2011) and has only two reported mutant alleles, *alc* and *nor* in tomato (Casals et al. 2012; Dias et al. 2003; Garg et al. 2008a). In this study, we identified a novel allele in Penjar-1 with two mutations, one with Q13K and second, where a stop codon terminated the NOR protein after six aa, while the other three Penjar accessions had *alc* mutation. All four accessions showed extended fruit shelf-life during post-harvest storage, a phenotype consistent with reported *nor*/*alc* mutants (Casals et al. 2012). Considering that post-harvest shelf life of Penjars is dependent on genetic background, the same may have contributed to the observed differences in the shelf life of these accessions (Casals et al. 2012; Garg et al. 2008a,b). The 55 days delay in SEN compared to AC strongly indicates that post-harvest shelf life is considerably prolonged in the Penjar fruits.

In *nor/alc* mutant, the delayed onset or total loss of ripening signifies that the *NOR* gene regulates a majority of the ripening triggered processes (Casals et al. 2015; Osorio et al. 2011; Saladié et al. 2007). Interestingly, despite mutations in *NOR* gene, the *RIN* expression was not considerably altered, except in Penjar-1, a total knockout mutant. Such a downregulation of *RIN* expression was also observed in *nor* and *SlNAC4* RNAi lines (Martel, 2010; Zhu et al. 2014). Evidently, though *RIN* is a master regulator, *NOR* acts autonomously of *RIN*, and optimal regulation of ripening requires a concerted action between these two regulators (Osorio et al. 2011).

A factor influencing fruit firmness is the cuticle as it prevents water loss and sustains cellular turgidity (Saladié et al. 2007). Consistent with earlier studies (Kosma et al. 2010; Saladié et al. 2007) the overall cuticle composition (wax and cutin) of Penjars is more similar to *nor/alc* mutants than AC. At the same time, the four Penjars widely vary in levels of different constituents of the cuticle. Only at MG, the Penjars share upregulation of few cuticle constituents and gene transcripts, whereas, at RR, such a shared regulation is largely absent. Surprisingly, post-MG despite variations in building blocks of the cuticle, all Penjars share prolonged shelf life and reduced water loss. Seemingly along with the composition, the cuticle architecture is also a key determinant of the prolonged shelf life of Penjars. The endogenous factors regulating cuticle composition/architecture are currently not known (Yeats and Rose, 2013; Fernández et al. 2016). It appears that the cuticle composition/architecture is not the sole factor, and the regulation of shelf life also involves other cellular processes altered by *NOR* mutation in Penjars.

In addition to the cuticle, the cell wall is also considered a key determinant for retention of fruit firmness. Consistent with this, the silencing of several cell wall modifying enzymes in tomato extends the shelf life of fruits (Vicente et al. 2007). The prolonged post-harvest shelf life indicates that the process of cell wall dissolution is considerably slower in Penjar fruits. The presence of several NAC domains in the promoters of cell wall modifying genes suggests that mutations in *NOR* would lead to lowering of their expression. Consistent with this, the expression of most cell wall modifying genes is downregulated during ripening in Penjars similar to *nor/alc* mutants (Osorio et al. 2011; Saladié et al. 2007). Even at SEN, several of these genes showed reduced expression indicating slower degradation of the cell wall, which is also corroborated by lower levels of cell wall related metabolites in Penjars than in AC.

One of the characteristic features of *nor/alc* fruits is the reduced accumulation of carotenoids compared to normal ripening cultivars (Giovannoni, 2007; Kopeliovitch et al. 1979; Sink Jr. et al. 1974). In Penjars, the reduction in carotenoids was manifested for most fruit-specific carotenoids including lycopene and β-carotene as well as for precursors, phytoene, and phytofluene. The reduced level of precursors is consistent with the lower expression of *PSY1* and *CYCB* in the Penjar fruits during ripening. In tomato fruits, carotenoid accumulation is also strongly influenced by ethylene; consequently *Nr* mutant of tomato shows diminished accumulation of carotenoids, due to a defect in ethylene perception (Barry et al. 2005). Similar to carotenoids, the reduction in ethylene emission from Penjars fruits was associated with lower expression of system II ethylene biosynthesis genes, *ACS2*, *ACS4* and *ACO1* (Barry and Giovannoni, 2007; Pech et al. 2012). The promoter analysis of carotenogenesis and ethylene biosynthesis genes revealed multiple NAC binding sites (Table S4) suggesting the downregulation of above genes may be related to the absence of a functional NOR protein in Penjars.

Current evidence indicate that fruit development is regulated by extensive interactions among different plant hormones (Kumar et al. 2014; Liu et al. 2015, Bodanapu et al. 2016). However, the overall hierarchical order of hormonal regulation in tomato development is not fully known. Hormonal profiling of Penjar fruits revealed that the *NOR* mutations influenced several hormones across the different developmental stages of fruits. In tomato, several studies have examined the influence of individual hormones on the accumulation of carotenoids (Kumar et al. 2014; Liu et al. 2015). However, a comprehensive correlation between hormones and carotenoids is largely missing. Barring Penjar-4, the interactions between hormones and carotenoids were similar in other Penjars albeit with few differences. The similarity in hormone-carotenoids interactions indicates that though *NOR* mutation attenuates carotenoid accumulation in Penjars, it does not influence the process underlying above interactions. The different interaction profiles in Penjar-4 may owe to nearly steady JA levels at all ripening stages. Evidence from JA-deficient mutants has implied that JA influences carotenogenesis in tomato independent of ethylene (Liu et al. 2012). The hormone-carotenoid interactions also indicate that in addition to JA and ethylene, other hormones also influence the regulation of carotenogenesis in tomato.

In tomato, suppression of ethylene biosynthesis delays ripening of fruits and extends the shelf life (Oeller et al. 1991). Consistent with this, lower ethylene emission may have contributed to the prolonged shelf life of Penjar fruits. Also, lower ABA levels at MG and RR in Penjars can lead to slower ripening (Sun et al. 2012). The accumulation of ABA is also characteristic of plant cells that retain water (Iuchi et al. 2000; Wan and Li, 2006). The higher retention of water in Penjar fruits may be assisted by higher ABA levels observed at SEN in all Penjars. Consistent with this view, higher sugar levels in Penjars may help in retention of cellular turgidity as marked by slow water loss (Vicente et al. 2007). A similar correlation between sucrose levels and increased firmness was also observed in *LIN5* suppressed fruits (Vallarino et al. 2017).

The extension of shelf life of fruits demands an optimal utilization of stored resources. During post-harvest storage, the fruits are deprived of support from the mother plant and can stay fresh only by lowering metabolism. Consistent with this, Penjar fruits show reduced metabolic processes such as turnover of proteins, fatty acids and reduced flux via TCA cycle. The lowering of citrate and upregulation of malate indicates an attenuation of respiration that is critical for prolonging the shelf life (Centeno et al. 2011). Likewise, abundances of the aspartate amino acid family (methionine, isoleucine, threonine, and lysine) have been linked to energy metabolism during seed germination (Angelovici et al. 2011; Kirma et al. 2012). The downregulation of isoleucine, threonine observed in Penjars at SEN indicates reduced flux in TCA cycle. Increased lysine levels in Penjar-2 and -4 in SEN fruits was similar to that found in the fruits of *LeACS2* suppressed transgenic line (Gao et al. 2007). Lowering of metabolism was not restricted to respiration, but even protein turnover was reduced, as evident by the decrease in the levels of free amino acids in Penjars. The lowering of metabolism also requires a balance between carbon and nitrogen metabolism, which is consistent with a reduction in GABA levels in Penjars, as GABA acts as a regulatory molecule to fine tune the cooperation between these two pathways (Takayama and Ezura, 2015). Collectively, above metabolic shifts indicate attenuation of overall metabolism during ripening and post-harvest storage of Penjars, contributing to their long shelf life.

In summary, our study revealed a wide-ranging influence of *NOR* mutations on diverse metabolic processes in four different Penjars. The prolonged shelf life of Penjars appears to involve attenuation of several metabolic processes associated with slower degradation of cell walls. The sustenance of firmness seems to be correlated with higher sucrose and reduced water loss, and hitherto unknown features of cuticle composition/architecture in the Penjars. In future, a better comprehension of the metabolic processes including an understanding of cuticle architecture combined with genome editing of causative genes would facilitate the improvement of the shelf life of perishable fruits.

## Acknowledgements

We thank Department of Biotechnology, Government of India (BT/PR11671/PBD/16/828/2008) funding to YS and RS, Council of Scientific and Industrial Research fellowship, New Delhi to RK and DST-Fund for Improvement of Science and Technology to Department of Plant Sciences. We thank Rafael Fernandez Muñoz, Institute for Mediterranean and Subtropical Horticulture, Malaga, Spain for the kind gift of Penjar accessions.

## Author contributions

The work was conceptualized by RK and YS. Most experiments were done by RK and VT helped in the wax analysis. RK, RS, and YS were involved in writing the manuscript, and all authors read and approved the manuscript.

## Conflict of Interest

The authors declare no conflict of interest.

## Supplementary data

The following materials are available in the online version of this article.

**Figure S1.** Sequence analysis of *NAC-NOR* in Penjars.

**Figure S2.** Fruit phenotypes (A) and chronological development and ripening (B) in AC and Penjars.

**Figure S3.** The on-vine shelf life of Penjar and AC fruits.

**Figure S4.** Relative expression of *RIN* and *NOR* genes in AC and Penjar fruits.

**Figure S5.** Loss of fruit weight in both AC and Penjars after chloroform treatment (A) and the percent relative abundance of cuticular wax components in MG and RR fruits of AC and Penjars.

**Figure S6.** Relative expression of genes associated with cuticle biosynthesis in AC and Penjar fruits.

**Figure S7.** Location of putative NAC motifs in the promoters of cell wall modifying genes. **Figure S8.** Ethylene emission and relative expression of ethylene biosynthetic genes in AC and Penjar fruits.

**Figure S9.** Carotenoid composition in AC and Penjar fruits during ripening and post-harvest storage.

**Figure S10.** Relative expression of key carotenoid biosynthesis genes in AC and Penjar fruits.

**Figure S11.** Correlation network of carotenoids, carotenogenic genes, and hormones during fruit ripening.

**Figure S12.** Principle component analysis (PCA) of metabolic profiles in AC and Penjar fruits.

**Figure S13.** The relative abundances of organic acids in the AC and Penjar fruits during ripening and post-harvest storage.

**Figure S14.** The relative abundances of amino acids and amines in the AC and Penjar fruits during ripening and post-harvest storage.

**Figure S15.** The relative abundances of sugars, sugar alcohols and sugar-derived acids in the AC and Penjar fruits during ripening and post-harvest storage.

**Figure S16.** The relative abundances of miscellaneous compounds (fatty acids, vitamins and vitamin-derivatives, nucleotides, phenolics, etc.) in the AC and Penjar fruits during ripening and post-harvest storage.

**Table S1.** Primers used for PCR in the present study.

**Table S2.** Carotenoid content in AC and Penjar fruits.

**Table S3.** The interactions between different metabolites and genes in the correlation networks of AC and Penjar fruits.

**Table S4.** List of NAC binding sites identified in the promoters of various genes.

**Table S5.** List of metabolites identified and their relative abundances in AC and Penjar fruits during ripening and post-harvest storage using GC-MS.

**Table S6.** List of cuticular wax components identified and their relative abundances in AC and Penjars during ripening using GC-MS.

